# Internal DNA standards for Isopycnic Centrifugation of Environmental DNA in Quantitative Stable Isotope Probing

**DOI:** 10.1101/2025.06.27.661829

**Authors:** Egbert Schwartz, Michaela Hayer, Rebecca L. Mau, Katheryn Nantz, Bruce A. Hungate, Steve Blazewicz, George Allen, Ember Morrisey, Binu Tripathi, Jennifer Pett-Ridge

**Author notes:** Corresponding Author: Egbert Schwartz –.

## Abstract

Element assimilation rates or the DNA replication rate of microbial taxa can be measured in environmental samples through quantitative DNA-stable isotope probing (DNA-qSIP). Here, we introduce a set of DNA standards that may be used to quantify the density of DNA extracted from environmental samples after isopycnic centrifugation. The standards are approximately 9 Kbp PCR products with either isotopically enriched (98 atom% ^15^N, ^13^C) nucleotides or natural abundance nucleotides and have densities that differ by 0.051 g/mL. The internal standards were tested in a DNA-qSIP analysis of bacterial populations in soil exposed to 63 atom% H_2_^18^O, performed in two different laboratories with different equipment and protocols. While fractionation results, including number of fractions taken and differences in density between adjacent fractions, between the two laboratories were different, the internal standards allowed the two data sets to be compared, and both research groups found similar Excess Atom Fraction (EAF) of oxygen-18 in the DNA of bacterial taxa. These internal DNA standards allow direct comparison of DNA-qSIP results from different experiments regardless of operator, tracer enrichment levels, protocol or equipment used and can support the creation of a large global database that contains qSIP results from many different laboratories.

**Importance:** DNA-qSIP is an important technique in microbial ecology that allows the growth and nutrient assimilation rates of microbial taxa, including those that have not yet been cultured, to be measured. Growth and assimilation rates are extremely variable parameters in microbial ecology because most of the microbial community is dormant and inactive, which complicates linking microbial populations to ecosystem processes. DNA-qSIP is increasingly practiced in a range of laboratories, using different equipment, isotopes, substrates, and protocols, making comparison of results among different research groups challenging. Here we describe the development of internal standards, 2 pieces of DNA with different isotopic content and hence buoyant density, that can be added to environmental DNA before isopycnic centrifugation. Our results show the internal DNA standards allow DNA-qSIP results to be compared across laboratories and could contribute to the formation of a large DNA-qSIP database that contains growth or nutrient assimilation rates of microbial taxa from many different experiments.

## Introduction

With DNA quantitative Stable Isotope Probing (DNA-qSIP) experiments, the assimilation of a heavy isotope, such as oxygen-18, into microbial genomes within an environment is quantified (Blazewicz et al., 2020; Hungate et al., 2015). The technique requires comparison between DNA extracted from microbial communities exposed to a substrate with a high EAF of the rare heavy isotope and DNA from samples exposed to natural abundance levels of heavier isotopes. The DNA is separated along a cesium chloride density gradient formed during isopycnic centrifugation in an ultracentrifuge. The contents of the DNA in the ultracentrifuge tube are collected in many fractions each with a slightly different density. By analyzing the DNA in each of the fractions through sequencing and qPCR it is feasible to determine the EAF of microbial genomes which in turn, may be related to microbial growth or nutrient assimilation rates.

Many laboratories across the globe conduct DNA-qSIP experiments, but it remains difficult to directly compare results between experiments. The precision and accuracy of DNA-qSIP results can vary widely among experiments and the impact of different operators, centrifugation protocols, equipment and qualities of environmental DNA extracts on the EAF measurements made during DNA-qSIP remains unknown. There have been many differences in experimental protocols among DNA-qSIP experiments. Both vertical (Blazewicz et al., 2020; Nuccio et al., 2022) and side angle rotors (Bell et al., 2023; Hayer et al., 2022; Hungate et al., 2015) have been used. The volume of ultracentrifuge tubes and centrifugation protocols including g-force, and length of spin are different among published DNA-qSIP experiments. The level of isotopic enrichment during experimental incubations can also vary considerably and different isotopes including nitrogen-15, carbon-13 and oxygen-18, may be used in DNA-SIP experiments (Buckley et al., 2007; Radajewski et al., 2000; Schwartz, 2007).

Many studies of microbial communities have advocated the use of internal DNA standards to control for different DNA extraction efficiencies between samples (Crossette et al., 2021; Satinsky et al., 2013; Smets et al., 2016). These internal standards can consist of plasmid DNA or genomic DNA of organisms that are unlikely to be encountered in the environmental samples under investigation yet presumed to have an extraction efficiency equal to the DNA of native organisms. By quantifying the amount of extracted DNA standard through quantitative PCR or sequence read counts and comparing this quantity to the known amount of DNA standard added prior to DNA extraction, it is possible to calculate an extraction efficiency and determine the amount of DNA present in an environmental sample.

To date, DNA standards have not been commonly used in DNA-qSIP experiments to control for variations in isopycnic centrifugation, although Vyshenska and colleaugues (Vyshenska et al., 2023) added six different synthetic DNA fragments to environmental DNA prior to centrifugation to monitor the quality of density separations. In their study, synthetic DNA, approximately 2 Kbp in length with GC contents ranging between 37% and 63%, and with different ^13^C contents, were used so that they would separate along a cesium chloride concentration gradient in a predictable order. Ten ng of synthetic DNA were added to 1 µg of microbial DNA mixture. If the spike-in distribution patterns did not match the expected order along the density based on the theoretical estimated density of the spike-in (given its GC content and ^13^C/^12^C ratio), then the sample was considered problematic and removed from the analysis. However, no analytical approach was described to compare samples from different research groups to each other.

Here we describe a set of internal DNA standards that can facilitate comparison of qSIP results among different laboratories by controlling for differences in protocols between DNA-qSIP experiments and allow all DNA-qSIP experiments to be compared to each other.

## Materials and Methods

### Construction of Internal Standards

Two types of plasmids, which served as templates for producing either Internal Standard 2 (IS2) or Internal Standard 3 (IS3), were constructed. The DNA primers used to synthesize the standards are listed in Table 1. Both consisted of the TOPO-2XL plasmid (Thermo Fisher Scientific, Waltham MA, USA) with a 5,026 bp insert constructed from *E. coli* genomic sequences (Figure 1). The inserted sequence contained conserved primer binding sites (515F, GTGCCAGCAGCCGCGGTAA) and (806R, ATTAGATACCCTGGTAGTCC) which are commonly used in sequencing a fragment of the 16S rRNA genes of bacterial communities (Apprill et al., 2015; Parada et al., 2016). Sequences taken 8,260 bp upstream (215,510 – 215,803 for IS2) or 7,569 bp upstream (216,201-216,490 for IS3) from the first 16S rRNA gene of E. coli, strain K12 were located in between the 2 sequencing primer sites so that the internal standards could be detected through routine 16S rRNA gene sequencing analysis.

**Figure 1.**
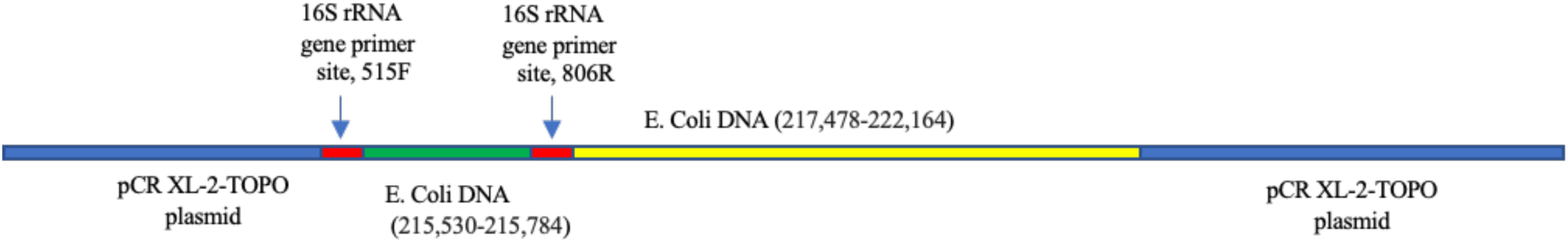
A map of Internal Standard 2, a PCR product generated from a plasmid template. The plasmid contains fragments of *E. coli* DNA taken upstream from one of the *E. coli*’s 16S rRNA genes and sites that can be recognized by standard bacterial 16S rRNA gene sequencing primers.

**Table 1.**
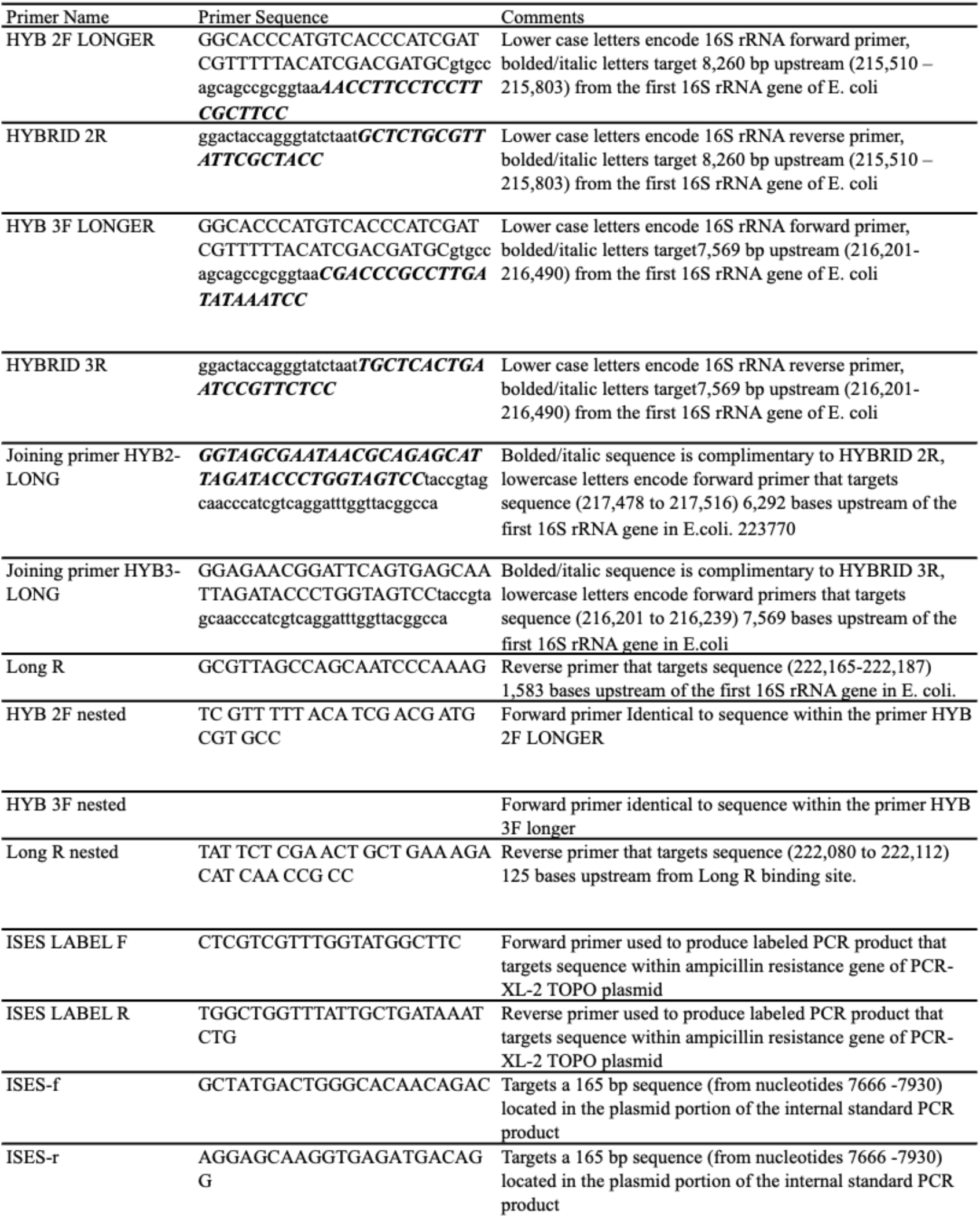
A description of DNA primers used to construct, label, and detect internal quantitative Stable Isotope Probing standards.

The sequence inserted into the plasmid was produced through a series of three PCRs (Supplemental Figure 1). In the first reaction, *E. coli* K12 genomic DNA was used as a template with the primers HYB 2F LONGER and HYBRID 2R for IS2 or HYB 3F LONGER and HYBRID 3R for IS3 (see Table 1 for primer sequences and template annealing sites). The ingredients for the PCR were as follows: 25 μL SuperFi master mix (Thermo Fisher Scientific, Waltham MA, USA), 14 μL H_2_O, 5 μL GC enhancer, 2.5 μL of each 20 μM primer and 1 μL of *E. coli* K12 genomic DNA (10 ng/µL) which was extracted with a Wizard genomic DNA purification kit (Promega inc., Madison WI, USA). The reaction was started with a hot start at 95°C for 5 min., followed by 30 cycles of 95°C for 10 sec., 60°C for 10 sec., and 72°C for 120 sec., followed by a final extension step at 72 °C for 5 min. In the second PCR a fragment of *E. coli* genomic DNA was amplified with either the primers Joining Primer HYB2-LONG and LONG R for IS2 or Joining Primer HYB3-LONG and LONG R for IS3. PCR products were cleaned to remove remaining primers and nucleotides with a PCR cleanup kit (Qiagen, Germantown MD, USA). The two PCR products have complimentary sequences and were joined together in a PCR without primers. The PCR contained the following ingredients: 1 µL of the HYB 2F LONGER and HYBRID 2R PCR product or 1 µL of the HYB 3F LONGER and HYBRID 3R PCR product, 15 µL of Joining primer HYB2-LONG and LONG R PCR product or Joining primer HYB3-LONG and LONG R PCR product, 25 µl Super Fi master mix, 5uL of GC enhancer, and 4 µL of H_2_O. The PCR program was as follows: a 30 sec. hot start at 95°C, 15 cycles of 10 sec at 95 °C, 10 sec. 60°C, and 120 sec. at 72°C, followed by a final 5 min. extension step at 72°C. The joined PCR product was used as a template in a third PCR. This PCR contained 2.5 µL of the primer HYB 2F nested (for IS2) or HYB 3F nested (for IS3), 2.5 µL of the primer Long R nested, 1 µL of the cleaned joined PCR product, and 14 µL of water. The PCR program was as follows: a 30 sec. hot start at 95°C, 27 cycles of 10 sec at 95 °C, 10 sec. 60°C, and 120 sec. at 72°C, followed by a final 5 min. extension step at 72°C. This PCR product was cleaned and then cloned into a PCR-XL-2 TOPO plasmid according to manufacturer instructions (Thermo Fisher Scientific, Waltham MA, USA).

Plasmids were extracted from cultures which contained either the IS2 or IS3 plasmids using a Qiagen Plasmid Extraction kit and were used as a template in a PCR with the primers IS LABEL R (TGGCTGGTTTATTGCTGATAAATCTG) or IS LABEL F (CTCGTCGTTTGGTATGGCTTC). The PCR with IS2 plasmid template contained either 98 atom% ^15^N and ^13^C nucleotides, producing the PCR product IS2L, or nucleotides with natural abundances of carbon and nitrogen isotopes, forming the PCR product ISR2. The reaction with IS3 plasmid as a template contained nucleotides with natural abundances of all isotopes and produced the PCR product ISR3. The ingredients in the PCR were as follows: 2.5 µL each of the primers ISES LABEL F and ISES LABEL R (both at 10µM), 1 µL of extracted plasmid, 29.5 µL of water, 10 µL of 5X GC buffer, 4µL of 2.5 mM dNTP’s and 0.5 µL of Phusion DNA polymerase. The PCR program was as follows: a 30 sec. hot start at 95°C, 27 cycles of 10 sec at 95 °C, 10 sec. 60°C, and 120 sec. at 72°C, followed by a final 5 min. extension step at 72°C. The isotopically labeled PCR product IS2L contained 26.5 more neutrons per basepair of DNA than IS2R or IS3R PCR products. All three PCR products were used as internal DNA standards in the following DNA-qSIP experiment.

The internal DNA standards were added to DNA extracted from a Basalt soil sample collected from the rhizosphere of a Blue Gramma plant in Flagstaff, AZ, USA at 35.21283° N, 111.65422° W. The DNA was extracted with a DNeasy PowerMax soil kit (Qiagen, Germantown MD). Three 10-gram replicates of the soil were incubated with 2.5 mL of 98 atom% ^18^O-water each, while another three 10-gram replicates were incubated with water containing natural abundance levels of oxygen-18. DNA was extracted after the soils were incubated at room temperature for eight days. The moisture content of the soil, after water addition, was 38.9% and the soil water from samples incubated with isotopically labeled water had an oxygen-18 content of 63 atom %.

DNA-qSIP with the internal DNA standards was performed in two different laboratories: the Laboratory for Isotopic and Molecular Ecosystem Science at Northern Arizona University (NAU) in Flagstaff, AZ and at Lawrence Livermore National Laboratory (LLNL) in Livermore, CA. We first describe the DNA-qSIP procedure that was used at NAU and then the one employed at LNLL.

To separate DNA by density, 5 μg of DNA was added to 3.6 mL of saturated cesium chloride solution and 200 µL of gradient buffer (0.2 M Tris, 0.2 M KCl, and 2 mM EDTA) in 4.7 ml OptiSeal polyallomer tubes (Beckman Coulter, Brea, CA, USA). The internal DNA standards IS2L and IS3R were added to the extracted soil DNA at three different concentrations: 1 ng internal standard per 1µg of soil DNA, 0.1 ng of Internal Standard per 1 µg of soil DNA, or .01 ng internal standard per 1 µg of soil DNA. After centrifugation at 127,000 x *g* at 18°C for 72 h in a Beckman TLA-110 rotor in an OptimaTM MAX ultracentrifuge (Beckman Coulter, Brea, CA), the gradient was separated into approximately 22 - 200 μL fractions using a fraction recovery system (Beckman Coulter, Brea, CA). The density of each fraction was measured with a Reichert Refractometer. Each fraction was cleaned through isopropanol precipitation and a 70% ethanol wash. The High Throughput DNA-qSIP procedure used at LLNL is described in Nuccio et al., 2022. It employs a different ultra centrifuge (Beckman Coulter Optima XE-90), rotor (VTi65.2), spin time (108 hours at 176,284 X g), fraction collection system, and DNA cleanup procedure, using Polyethylene Glycol, instead of isopropanol, to precipitate the DNA. Fraction density graphs, which relate the fraction number to its density were generated from this data.

### Detection of the Internal DNA Standards in fractionated DNA with qPCR

Triplicate qPCR reactions were run to quantify the concentration of the DNA standards across sample fractions. Each reaction contained 1μL DNA template, 0.25 μM of forward (ISES-f GCT ATG ACT GGG CAC AAC AGA C) and reverse (ISES-r AGG AGC AAG GTG AGA TGA CAG G) primers, 1X Forget-Me-Not EvaGreen qPCR Master Mix (Biotium, Fremont CA), and 1.5 mM MgCl_2_ (10 µL reactions). The PCR program was as follows: 95 °C for 2 min with 35 cycles of 95 °C for 30 sec., 62 °C for 10 sec., and 72 °C for 10 sec.

### Detection of Internal DNA Standards with Illumina sequencing targeting 16S rRNA genes

Because the internal standards contain the Earth Microbiome Project primer sites (Apprill et al., 2015; Parada et al., 2016), it is possible to detect them in Next-Generation Sequencing analysis of 16S rRNA genes. However, the sequence between the two primer sites is not present in any standard 16S rRNA database because it is generated from a non-16S rRNA sequence in the *E. coli* genome and is removed from data sets during the denoising bioinformatics step using the DADA2 (Callahan et al., 2016) plugin in QIIME2 (Bolyen et al., 2019). Therefore, the abundance of internal standards in each fraction from samples analyzed at LLNL was quantified by mapping raw sequencing reads against the IS2 standard using BBMap (bbmap.sh in1=A16f1_8_L001_R1_001.fastq in2=A16f1_8_L001_R2_001.fastq out=A16f1.sam ref=IS2.fa nodisk). The .sam files were converted to .bam format using samtools for extracting the mapping details (samtools view -S -b A16f1.sam > A16f1.bam) after which the number of mapped mated pairs were extracted (samtools view -f 3 A16f1.bam | wc -l #divide the output number by 2).

### 16S rRNA gene qPCR and sequencing

Standard curves for qPCR of 16S rRNA genes were generated using 10-fold serial dilutions of genomic *E. coli* DNA (ATTC, MG1655). The 10 μL reactions contained 0.2 μM of the primers EUB 338F/EUB 518R (Fierer et al., 2005), 1X Forget-Me-Not EvaGreen qPCR Master Mix, and 1.5 mM MgCl_2_. The assay was performed on a CFX 384 (Bio-Rad, Hercules, CA), using a program of 95 °C for 2 min with 35 cycles of 95 °C for 10 sec., 62 °C for 10 sec., and 72 °C for 10 sec. For sequencing, two PCR steps were used (Berry et al., 2011). Each sample was first amplified using primers 515F and 806R (Parada 2016, Apprill 2015). This was done in triplicate 10 µL PCR assays containing 1 μM of each primer, 1X Phusion Green Hot Start II HF Mastermix (Thermo Fisher Scientific), and 1.5 mM MgCl_2_. PCR conditions were 95°C for 2 min; 15 cycles of 95°C for 30 sec., 55°C for 30 sec., and 60°C for 30 sec. Initial PCR products were pooled, 10-fold diluted, and used as template in the subsequent reaction with region-specific primers that included the Illumina flow cell adapter sequences and 8 nucleotide Golay barcodes (10 cycles identical to initial amplification conditions). Products of the reaction were purified with carboxylated SeraMag Speed Beads (Sigma-Aldrich, St. Louis, MO) and quantified by Picogreen fluorescence. Equal quantities of the reaction products were pooled, bead-purified once again, and quantified by qPCR using the Library Quantification Kit for Illumina (Kapa Biosciences, Woburn, MA), sequenced on a MiSeq instrument (Illumina, San Diego, CA) using 2 x 150 paired-end read chemistry.

### Analysis of Internal DNA Standard Results

The calculations used to compare qSIP samples with the internal DNA standards are outlined in a spreadsheet in the supplemental materials. Most of an internal DNA standard was recovered in three adjacent fractions. To standardize the separation between internal standards in all samples, the fraction number was treated as a continuous variable, so that the relative abundance peaks of internal standards or 16S rRNA genes of individual taxa could be identified on a continuous scale. The relative abundances of internal DNA standards in the three fractions were used to calculate a weighted average fraction (WAF) for each internal standard where y_max_ is the relative abundance of internal DNA standard in the fraction with the highest relative abundance, y_-1_ is the relative abundance of internal DNA standard in the lighter fraction adjacent to y_max_, y_+1_ is the relative abundance of internal DNA standard in the heavier fraction adjacent to y_max_ and f refers to the fraction number, a whole integer.

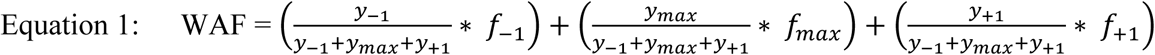

The difference between the WAF of the heavy (isotopically labeled) and the light (natural abundance) internal standards is defined as the span. Given differences in experimental protocols, spans may vary between labs and even within labs between samples. The span of all samples was adjusted to equal 1, a standard value in normalization procedures, by adding a span adjustment (*x_sample_*) to the first fraction. The *x_sample_* is different for each sample and equals:

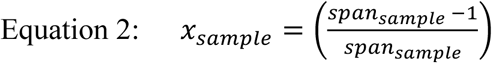

The adjustment resulted in new fraction numbers (*f_new_*) for each fraction, where n is the original fraction number.

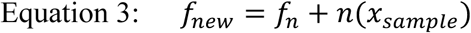

Subsequently, we adjusted the fraction numbers so that the peaks of the labeled internal DNA standard occurred at position 0 while the natural abundance internal DNA standard was positioned at 1 by adding a second adjustment to the new fraction number calculated above. The second adjustment equaled the WAF of the labeled internal standard peak, which was subtracted from each new fraction number, producing the final WAF that was used in subsequent calculations (see Supplemental Figure 2).

A linear trend line was fitted to a graph of the density of a fraction versus its WAF generated from the samples analyzed at LLNL and resulted in the following standard curve:

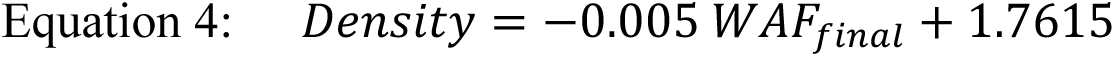

The linear fit had an R^2^ value of 0.998. Once a density was assigned to each fraction the data was run through the DNA-qSIP pipeline as described in (Hayer et al., 2022).

## Results

We created two different DNA molecules with identical sequences but different ^13^C and ^15^N contents to serve as internal standards during isopycnic centrifugation in qSIP analysis (Figure 1). The internal DNA standards are PCR products, approximately 9 Kbp long, and contain fragments of *E. coli* genomic DNA and the pCR XL-2-TOPO plasmid sequence. Primer sites complimentary to commonly used sequencing primers were engineered into the internal DNA standards so that they could be detected through standard DNA sequencing targeting the bacterial 16S rRNA gene.

There was a strong linear relationship between fraction number and the density of DNA as quantified with a refractometer (Figure 2A). Correlation coefficients ranged from 0.995 to 0.999. While these curves were all strongly linear, the slope and intercept varied, especially of samples processed at NAU. The linear fit for the fractionation curves obtained at NAU was density = -0.0066 * fraction number + 1.786. The standards deviations among the slopes and intercepts of these curves were 0.00026 and 0.0039, respectively. The linear fit for samples fractionated at LLNL was density = -0.0058 * fraction number + 1.788. The standard deviations for slope and intercepts were 0.00011 and 0.0018 respectively.

**Figure 2.**
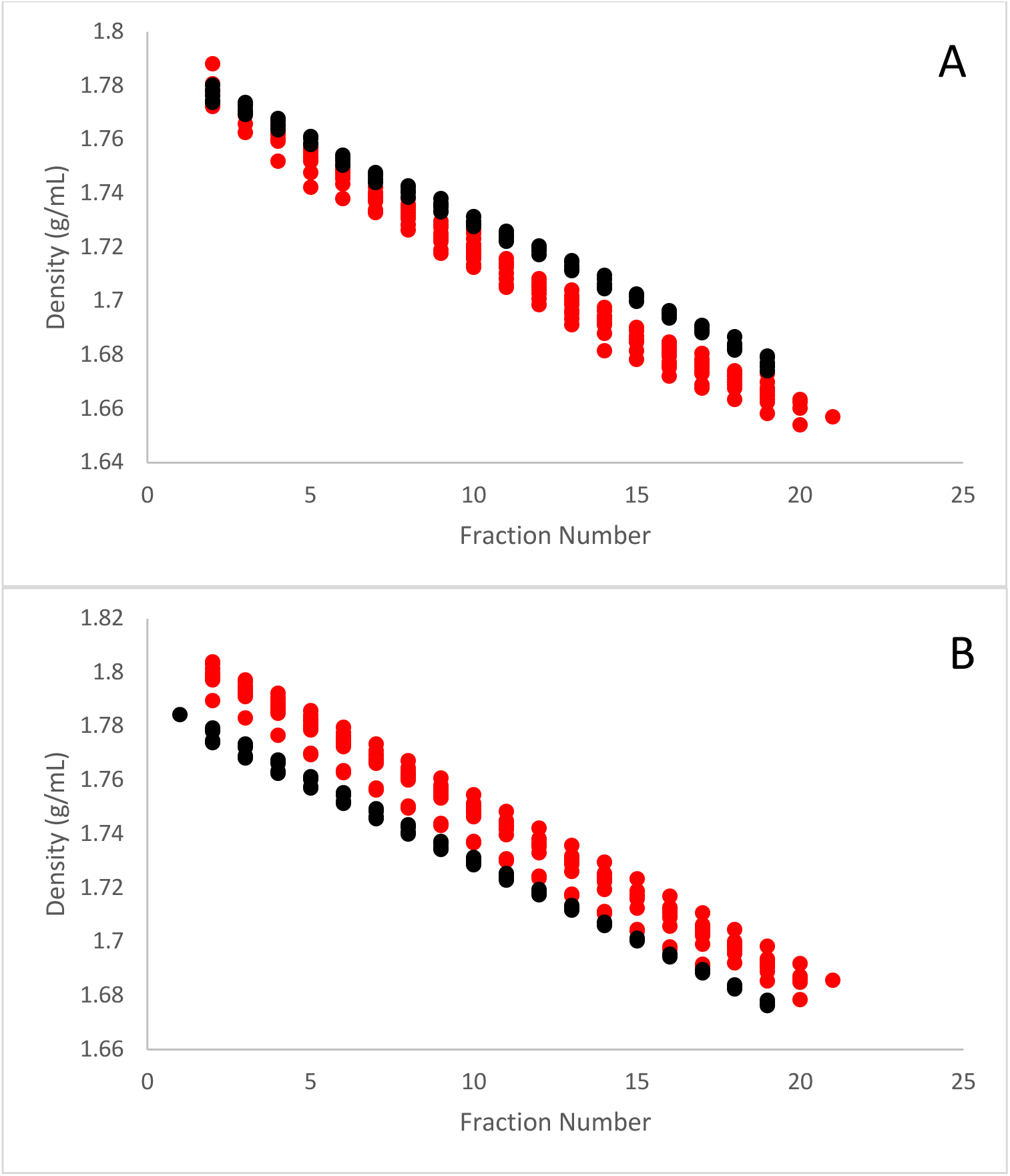
Comparison of fractionation curves obtained at Northern Arizona University (red) or Lawrence Livermore National Laboratory (black). The top graph shows fraction number versus densities determined with a refractometer while the bottom panel shows the relationship between fraction number and density measured with internal standards.

There was also a consistent linear relationship between fraction number and density calculated through internal DNA standards (Figure 2B). The linear fit of all the density curves generated at NAU with the internal standards was equal to density = -0.0064 * fraction number + 1.812 with standard deviations of 0.00018 and 0.0043 for the slope and intercept, respectively. In comparison, the linear fit of densities calculated with internal standards for samples processed at LLNL was density = -.0059 * fraction number + 1.789 and had standard deviations of 0.00015 and 0.0027 for the slope and intercept, respectively. The density of DNA in fractions varied from 1.68 to 1.78 g/mL.

Among different samples, the internal DNA standards consistently produced two distinct peaks in separate fractions reflective of the heavier (i.e. isotopically enriched) standard and the lighter (natural abundance) standard. However, these standards were not distributed identically in all samples, indicating that small differences in analytical conditions affected the gradient position of the internal DNA standards (Figure 3A). Most of each internal DNA standard could be recovered in 3 separate fractions, one fraction with a maximum amount of internal DNA standard copies flanked by two adjacent fractions. A WAF was calculated for both the labeled and natural abundance standards by weighting each of the three fractions by the part of total internal standard copies contained within that fraction. The WAF numbers in each sample were adjusted so that the light internal standard equaled a WAF of 1 and the heavier internal standard had a WAF of 0. By applying a standard curve generated at LLNL that related fraction number to density (equation # 4) it was possible to calculate the densities of fractions. Upon applying these normalization calculations, the internal standards exhibited near identical distributions across the density range (Figure 3B). After normalization, the internal DNA standards could be detected as two separate peaks with densities that differed by 0.051 g/mL. Once normalized densities were assigned to the fractions, it was possible to construct graphs relating fraction number to density (Figure 2B).

**Figure 3.**
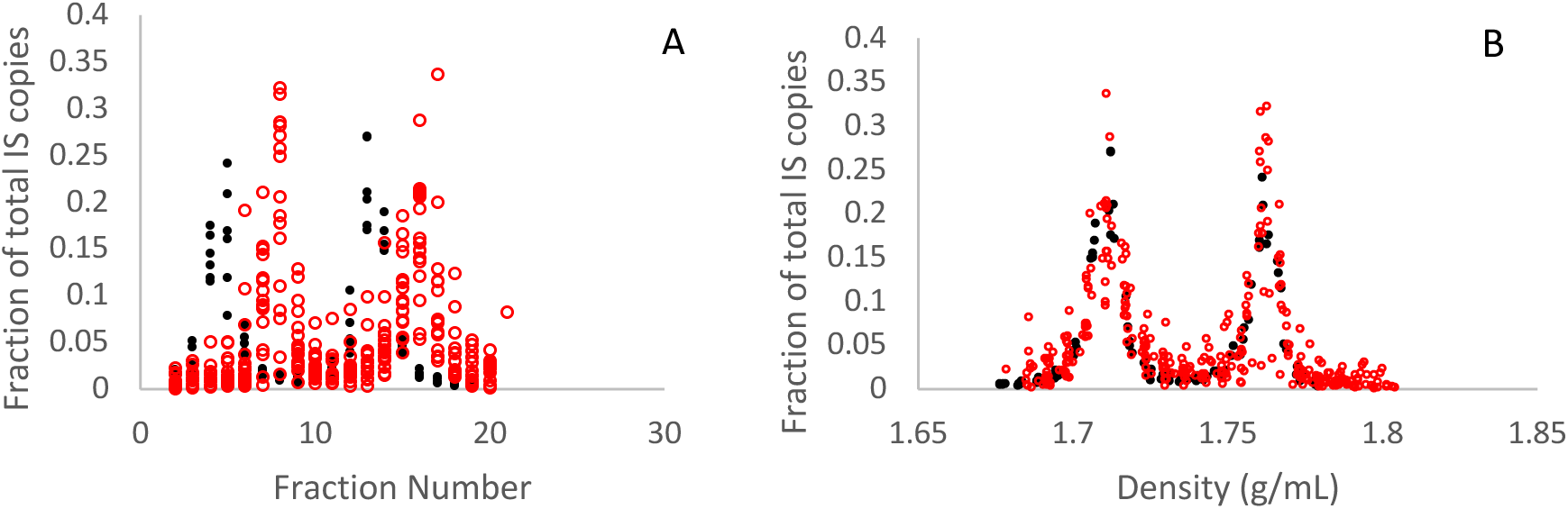
Comparison of internal standard distributions in raw (A) and transformed (B) data. Samples were analyzed at Northern Arizona University (○) or Lawrence Livermore National Laboratory (●).

The same DNA samples were centrifuged and fractionated in two different laboratories at NAU and at LLNL. There was greater variance in placement of the density curves generated at NAU, while the curves produced at LLNL showed a consistent relationship between fraction number and density (Figure 2). The slopes of the curves, but not the y intercepts, were more similar when the densities were determined with internal DNA standards than when the slopes were calculated using densities measured with a refractometer.

To assess the utility of internal DNA standards for normalizing densities, we examined the density distribution of DNA extracted from soil incubated with natural abundance ^18^O-water or 63 atom% ^18^O-water using densities derived from refractometry measurements (Figure 4a) and our internal standard approach (Figure 4b). Compared to DNA extracts from soils receiving natural abundance ^18^O-water, extracts from soils incubated with 63 atom % ^18^O-water had more DNA in fractions with densities greater than 1.73 g/mL. The method used to detect internal standards, either qPCR or sequence counts, did not have a large effect on the DNA-qSIP results (Figure 5). There was a strong positive correlation in the LLNL data set between the EAF’s of taxa measured with internal DNA standards detected through qPCR or sequencing (r^2^ = 0.997), though the slope of the linear trendline at 0.88 was less than 1 (Figure 5).

**Figure 4.**
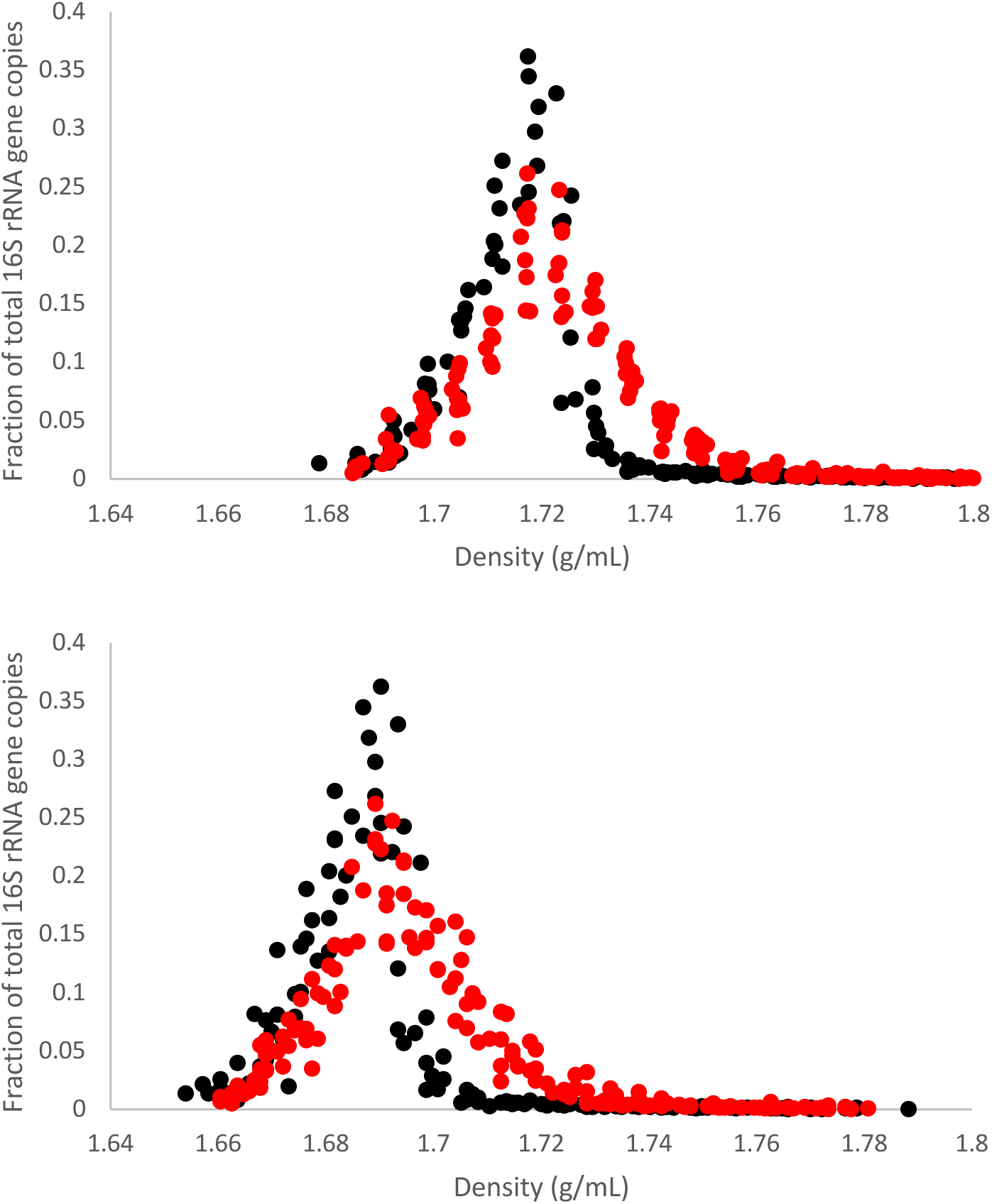
Comparison of DNA density curves from soil exposed to .63 Enriched Atom Fraction ^18^O-H_2_O (red) or .002 Enriched Atom Fraction ^18^O-H_2_O (black). The top panel shows the relationship with densities derived from refractometry measurements while the bottom graph used densities measured with the internal standards.

**Figure 5.**
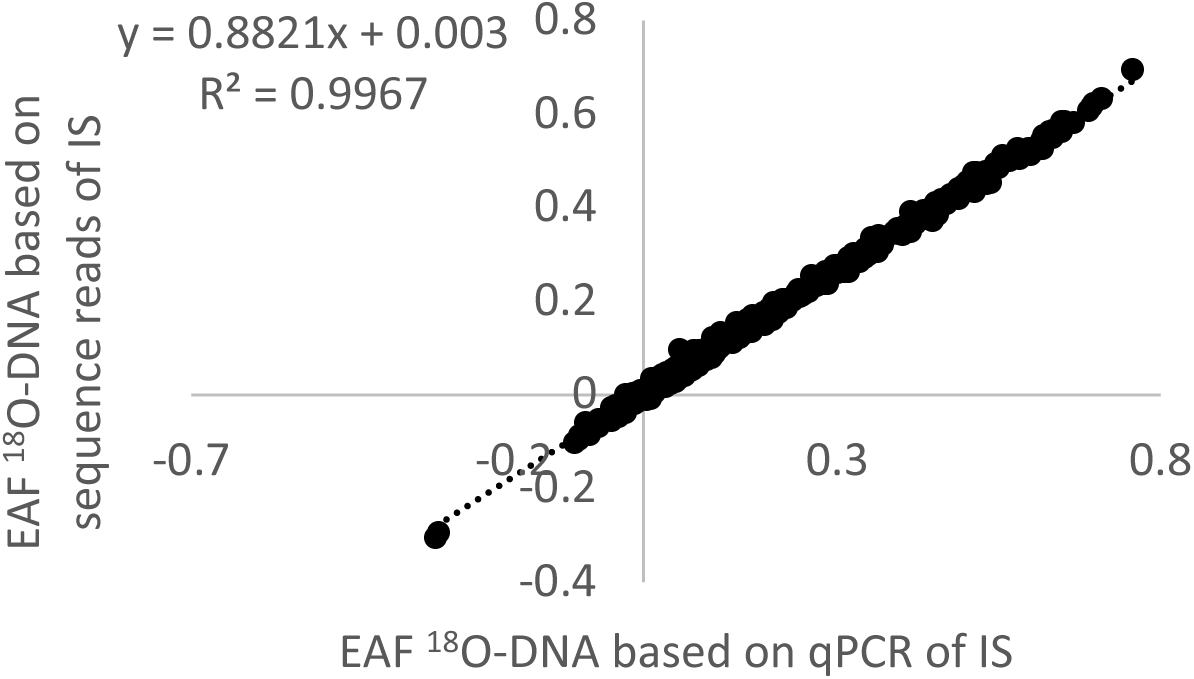
Comparison of Enriched Atom Fraction ^18^O-DNA of bacterial taxa determined by either quantifying internal standards abundances through qPCR or the relative number of internal standard sequence reads in libraries.

qPCR results of fractionated soil DNA, to which different amounts of internal standards were added (Supplemental Figure 3), indicate not more than 0.1 ng per µg environmental DNA of internal standard should be added to DNA extracts prior to centrifugation, because when 1 ng of internal standard per µg of environmental DNA was added (Supplemental Figure 3A), the heavy internal standards were much more abundant in the heavier fractions than 16S rRNA genes

For samples processed both at NAU and LLNL, EAF values agreed well when calculated for taxa that were present in at least three fractions in all replicates (Figure 6). A regression of these filtered results had an r^2^ of 0.88. However, when all taxa were retained in the analysis, including taxa present in fewer than three fractions, the correlation was not strong, yielding a relationship with an r^2^ of only 0.33. Application of these more stringent filtering criteria removed 396 out of 507 taxa, but the remaining 111 taxa still accounted for 81% of the total number of 16S rRNA gene sequences.

**Figure 6.**
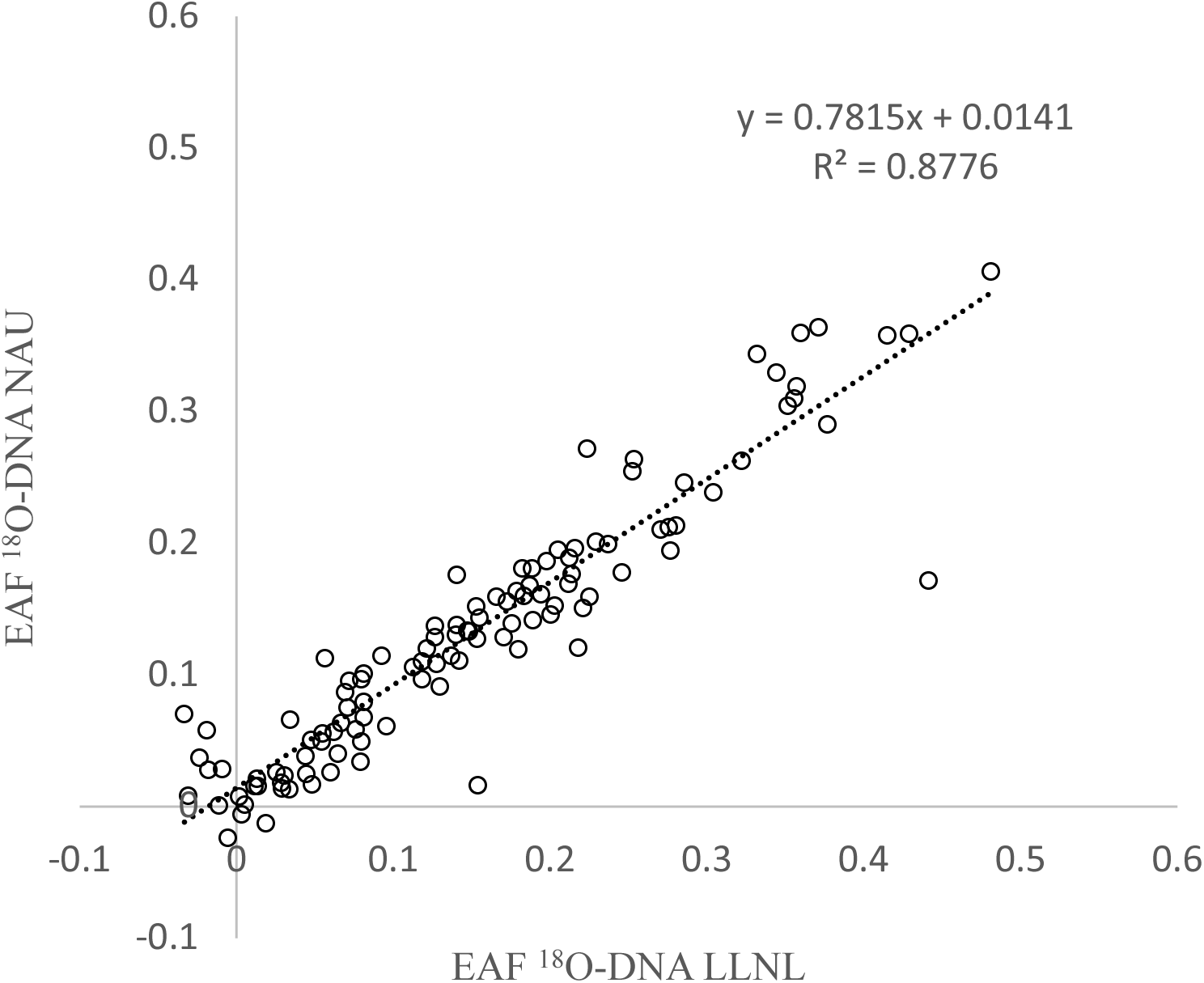
Comparison of Enriched Atom Fraction ^18^O-DNA of bacterial taxa measured at Northern Arizona University versus Enriched Atom Fraction ^18^O-DNA of bacterial taxa measured at Lawrence Livermore National Laboratory.

## Discussion

DNA-qSIP analysis, which measures the assimilation rates of isotopically labeled substrates into a microbial taxon’s genome, is increasingly used to study microbial population dynamics in environmental samples, (Blazewicz et al., 2020; Hungate et al., 2021). However, no single standard DNA-qSIP protocol exists, and research groups use different equipment and procedures, including different rotors, centrifuges, and fractionation equipment. Similarly, labs differ in the amount of DNA added for isopycnic centrifugation, cesium chloride concentrations, centrifugation parameters and DNA cleanup protocols. Furthermore, labs use different techniques (e.g., qPCR, sequencing, fluorimetry) to quantify DNA or specific sequences among the density gradient. Each of these methodological differences could contribute to lab-to-lab variation in EAF estimates, and a common internal standard could account for such differences.

As the number of DNA-qSIP studies increase there is an opportunity to establish a large DNA-qSIP database that allows comparison of a taxon’s growth or assimilation rates among many different studies. This database may be an important resource to identify and characterize microbial traits (Simpson et al., 2023). However, the variety of DNA-qSIP protocols may make direct comparison among experiments challenging. We propose that use of the internal DNA standards described in this manuscript will allow excess atom fraction of microbial taxa measured in different experiments to be related to each other.

The qSIP analysis performed at NAU included two different internal standards, IS2L and IS3R, which were identical in sequence except for the short sequence in between the two sequencing primer sites. As a result, these two internal standards can be distinguished from each other when the two internal standards overlap along the cesium chloride gradient and both standards appear in the same fraction. However, initial experiments showed that the labeled and natural abundance standards were not found in the same fraction and therefore it is not necessary to have two different internal standards. Because the relationship between fraction number and density (Figure 2A) is so strongly linear, we determined that only two internal standards are required to accurately characterize the density gradient.

We developed new internal DNA-qSIP standards by creating a plasmid that could serve as a template in a PCR with ^13^C and ^15^N enriched dNTP’s. We inserted additional *E. coli* DNA sequences into the plasmid to increase the size of the PCR product. Size affects the rate at which DNA sequences move to the position within the cesium chloride gradient that corresponds to their density (Meselson et al., 1957; Panijpan, 1977; Birnie & Rickwood, 1978). Increasing the size of the internal DNA standard ensured that the two standards would be fully separated after 72 hours of centrifugation.

The internal DNA standards can be added to the environmental DNA extracts prior to isopycnic centrifugation at very low concentrations (Supplemental Figure 3). This allows many research groups to share a single batch of internal DNA standards, ensuring that DNA-qSIP results can be compared among different laboratories within the larger scientific community. Only small amounts, less than 0.1 ng per μg environmental DNA, of internal DNA standards per ultra centrifuge tube are required--meaning it is feasible to make, in a single batch, sufficient internal DNA standards to analyze thousands of qSIP samples. The community of scientists engaged in qSIP could consider appointing a single organization to produce internal DNA standards so that all scientists can use the exact same preparation thereby reducing the variance among qSIP experimental results between different research groups.

The quantity of internal DNA standards added to a DNA-qSIP sample should be dependent on the method used to detect the internal standards in fractionated DNA. Very low amounts of internal standards, less than the 0.01 ng/µg soil DNA used in this study, are required if qPCR is used. The sequencing depth of a fraction’s DNA will determine the minimum of internal standards that can be detected through sequencing. Less environmental DNA is usually present in the denser fractions where the labeled DNA standard is found. In our studies, the labeled DNA standard was on average detected 4959 times in each fraction while the DNA standard made with natural abundance ^13^C and ^15^N isotopes was counted 256 times, indicating that nineteen times less labeled DNA standard could have been added to the samples for both DNA standards to be detected at similar rates (Supplemental Figure 4).

When comparing the raw data of DNA-qSIP results between the two different laboratories it was apparent that the internal DNA standards are not consistently recovered in the same fractions and that the data needs to be normalized so that the different samples can be compared to each other (Figure 3). There are two types of errors that needed to be corrected (Supplemental Figure 2). First, among samples the gradient positions of the two internal DNA standards were separated by different number of fractions. This could have been caused by centrifuging the samples at slightly different g forces. Faster spins result in a greater range of densities along the cesium chloride gradient and concentrate DNA into a tighter band that is recovered in fewer fractions. We refer to the number of fractions that separate the internal DNA standards as the span, and we adjusted the span for all samples to 1. The adjusted communal span could be, *a priori*, set to any number but 1 was picked because it facilitated easier calculations. The second error that needed to be standardized was that the gradient position of the internal DNA standards varied among samples. The position of a DNA molecule along a cesium chloride gradient is affected by the speed at which the sample is centrifuged but may also be impacted by the concentration of cesium chloride added to the sample at the beginning of the spin. We defined the gradient position of the labeled internal DNA standard to, a priori, be 0 and the gradient position of the natural abundance DNA standard to be 1, resulting in a span of 1. Again, these numbers were simply selected to facilitate easier calculations. The gradient positions of the internal DNA standards were adjusted *in silico* so that all DNA standards appeared at the same gradient positions in subsequent calculations.

Besides addressing the errors caused by using a refractometer due to evaporation of water from the samples, inaccurate cesium chloride standards or changes in temperature, the inclusion of internal DNA standards may eliminate the effort of making refraction measurements. Currently, these measurements must be made immediately after fractionating the cesium density gradient before the DNA is purified (Neufeld, Dumont, et al., 2007; Neufeld, Vohra, et al., 2007). Fractionation is a time-consuming effort, whether the process is automated or not, and not having to make refraction measurements should allow for more samples to be processed faster. Furthermore in some SIP studies, measurements of buoyant density using the refractometric index or by weighing are not always obtained (Simpson et al., 2023), nor is it always accurate due to inadequate refractometer calibration or challenges with weighing small volumes at high precision.

The DNA standards described in this paper can be detected in 16S rRNA amplicon sequencing because EMP primer binding sites (Caporaso et al., 2012) were incorporated into the internal DNA standards (Figure 1). Once the internal DNA standards are detected, the densities of the fractions, and subsequently, the EAF of bacterial taxa can be calculated. Some scientists may want to study other microorganisms, such as fungi or protists, through amplicon sequencing in qSIP experiments and may not intend to sequence the bacterial 16S rRNA gene. In these circumstances, it would be necessary to make new DNA standards which have the new sequencing primer sites that target the other microbial taxa, if the researchers intend to detect the internal DNA standards through sequencing. No modifications of the internal DNA standards are required if qPCR is used to quantify the abundance of the internal DNA standards in qSIP fractions. New internal DNA standards can readily be produced by following the instructions for making the internal standards outlined in the materials and methods section but using different HYB2F LONGER and HYBRID 2R primers. The new modified primers should have the lower-case sequences of these primers shown in Table 1 replaced with the sequence of the primers used to target the other microbial taxa. If shotgun sequencing is used, the number of reads that detect any sequence within the internal standards can be used to determine which fractions contain the internal standards.

Our comparison of qSIP data sets generated from the same environmental samples but produced by different laboratories showed that filtering out taxa present in fewer than 3 fractions is an important component of qSIP analysis. Without this filtering step there was much less agreement between the results produced in the 2 laboratories. EAF measurements of the genomes in rare taxa are likely less robust because the weighted average density of a taxa’s genome is determined from few fractions or data points. Large variations in EAF measurements can be observed in instances when taxa are present in only one natural abundance sample fraction and one enriched sample fraction. This problem can be addressed by sequencing the fractions more deeply so that rare taxa are detected in multiple fractions.

These internal DNA standards provide a method to calibrate DNA-qSIP data across experiments and laboratories. By assigning buoyant density without reliance on refractometry and enabling *in silico* normalization, this approach improves precision and enables direct comparison of microbial growth and assimilation measurements. This approach parallels standardization strategies used in other isotope-based fields, such as isotope-ratio mass spectrometry (Brand et al., 2014) and adopting such standards supports a more robust, interoperable and quantitative microbial ecology.

## Supporting information

Spreadsheet demonstrating Internal Standards calculation

## Acknowledgements

This work was supported by funding the U.S. Department of Energy, Office of Biological and Environmental Research, Genomic Science Program LLNL ‘Microbes Persist’ Soil Microbiome Scientific Focus Area (award #SCW1632). Work at LLNL was conducted under DOE Contract DE-AC52-07NA27344.

---

**Supplemental Figure 1.**
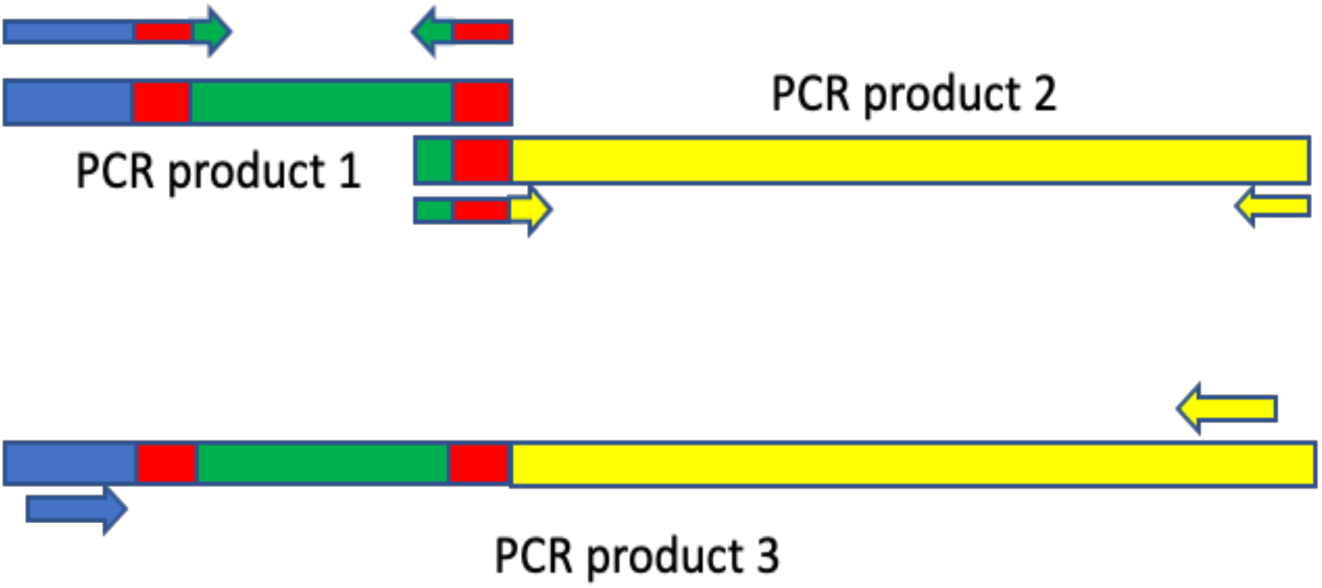
Illustration showing how the 3 PCR were used to construct a larger DNA sequence that was inserted into a plasmid. The plasmid was used in PCR to make internal standards with two different isotopic compositions. The Template for PCR 1 and PCR 2 was *E. coli* K12 genomic DNA while the two PCR products joined together served as a template for PCR 3.

**Supplemental Figure 2.**
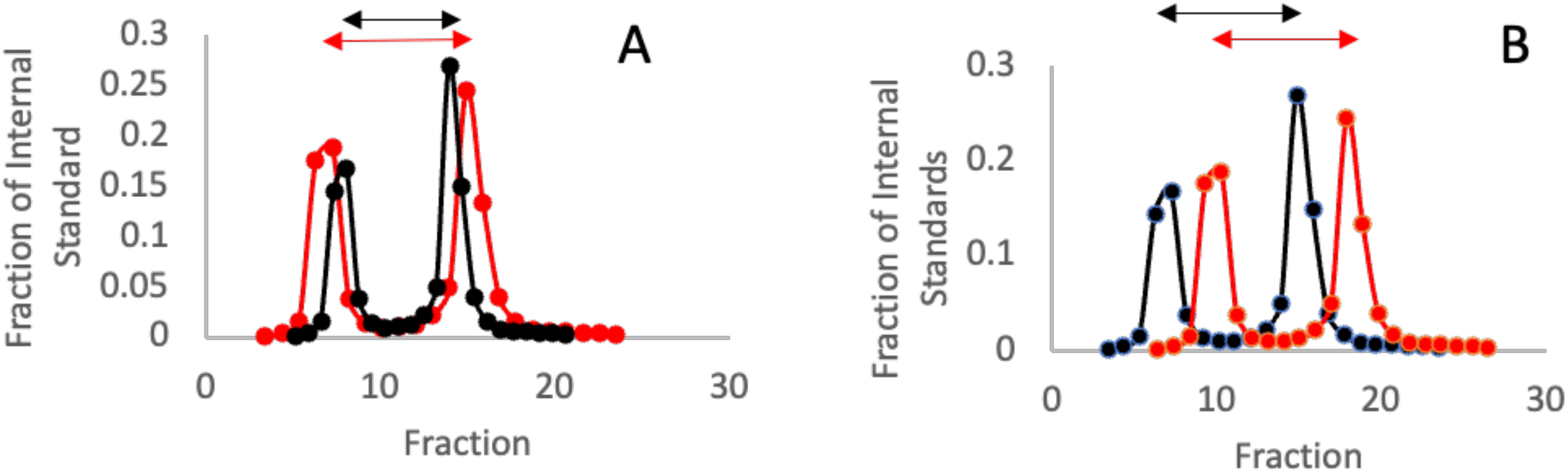
An illustration of the two types of adjustments required to compare results from 2 different ultra-centrifuge tubes using the internal standards. The two curves in each panel describe the proportion of an internal standard among fractions taken from the different centrifuge tubes. In Panel A the two standards are separated by a different number of fractions, the red arrow is longer than the black one. In panel B, the standards are separated by the same number of fractions but the position within the cesium chloride gradient is not the same. Here the red arrow is to the right of the black arrow.

**Supplemental Figure 3.**
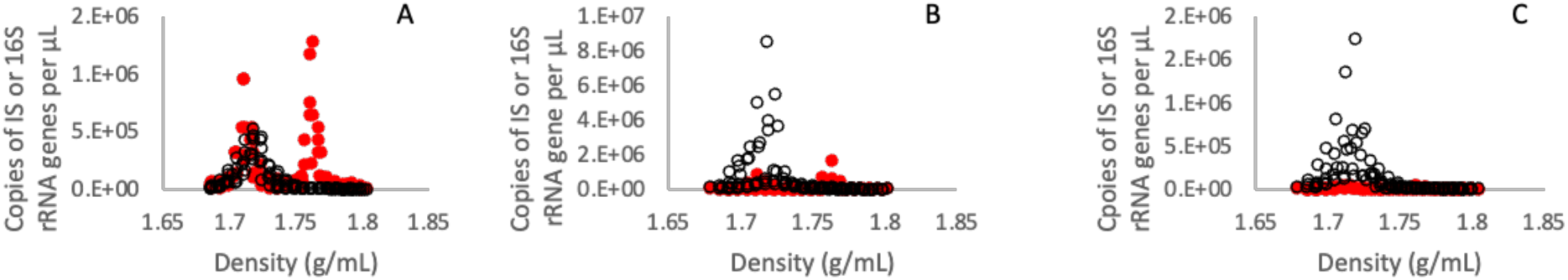
Comparison of the quantities of bacterial 16S rRNA gene copies (○) and DNA internal standards (●). A = 1ng DNA Internal Standard /µg of soil DNA, B = 0.1ng DNA Internal Standard /µg soil DNA, and C = .01 ng DNA Internal Standard /µg of soil DNA

**Supplemental Figure 4.**
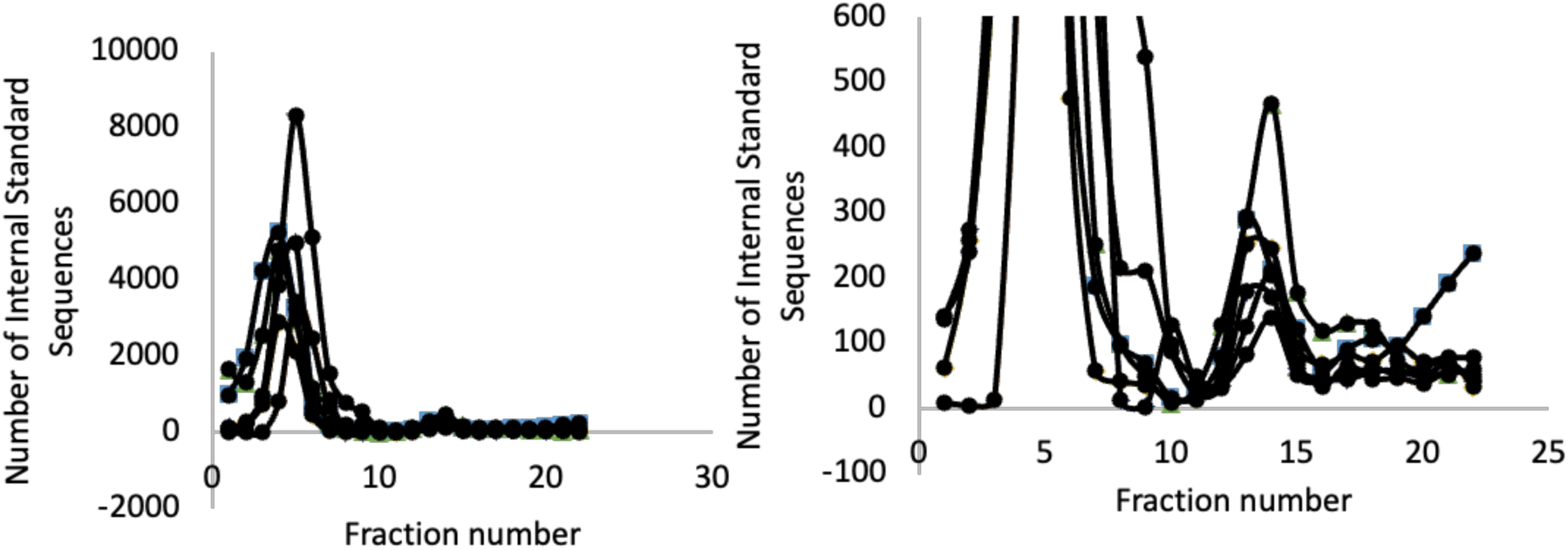
The number of DNA internal standard sequence reads in each fraction. Both panels show the same data, but the Y-axis scale has been adjusted between the two panels so that both internal standard peaks can be observed.

## References

Apprill, A., McNally, S., Parsons, R., & Weber, L. (2015). Minor revision to V4 region SSU rRNA 806R gene primer greatly increases detection of SAR11 bacterioplankton. Aquatic Microbial Ecology, 75(2), 129–137. 10.3354/ame01753

Aslett, D., Haas, J., & Hyman, M. (2011). Identification of tertiary butyl alcohol (TBA)-utilizing organisms in BioGAC reactors using 13C-DNA stable isotope probing. Biodegradation, 22(5), 961–972. 10.1007/s10532-011-9455-3

Bell, S. L., Zimmerman, A. E., Stone, B. W., Chang, C. H., Blumer, M., Renslow, R. S., Propster, J. R., Hayer, M., Schwartz, E., Hungate, B. A., & Hofmockel, K. S. (2023). Effects of warming on bacterial growth rates in a peat soil under ambient and elevated CO2. Soil Biology and Biochemistry, 178, 108933. 10.1016/j.soilbio.2022.108933

Berry, D., Mahfoudh, K. Ben, Wagner, M., & Loy, A. (2011). Barcoded primers used in multiplex amplicon pyrosequencing bias amplification. Applied and Environmental Microbiology, 77(21), 7846–7849.

Birnie, G. D., & Rickwood, D. (1978). Centrifugal separations in molecular and cell biology. Butterworths.

Blazewicz, S. J., Hungate, B. A., Koch, B. J., Nuccio, E. E., Morrissey, E., Brodie, E. L., Schwartz, E., Pett-Ridge, J., & Firestone, M. K. (2020). Taxon-specific microbial growth and mortality patterns reveal distinct temporal population responses to rewetting in a California grassland soil. The ISME Journal, 14(6), 1520–1532. 10.1038/s41396-020-0617-3

Bolyen, E., Rideout, J. R., Dillon, M. R., Bokulich, N. A., Abnet, C. C., Al-Ghalith, G. A., Alexander, H., Alm, E. J., Arumugam, M., Asnicar, F., Bai, Y., Bisanz, J. E., Bittinger, K., Brejnrod, A., Brislawn, C. J., Brown, C. T., Callahan, B. J., Caraballo-Rodríguez, A. M., Chase, J., … Caporaso, J. G. (2019). Reproducible, interactive, scalable and extensible microbiome data science using QIIME 2. Nature Biotechnology, 37(8), 852– 857. 10.1038/s41587-019-0209-9

Brand, W. A., Coplen, T. B., Vogl, J., Rosner, M., & Prohaska, T. (2014). Assessment of international reference materials for isotope-ratio analysis (IUPAC Technical Report). Pure and Applied Chemistry, 86(3), 425–467. 10.1515/pac-2013-1023

Buckley, D. H., Huangyutitham, V., Hsu, S.-F., & Nelson, T. A. (2007). Stable isotope probing with 15N2 reveals novel noncultivated diazotrophs in soil. Applied and Environmental Microbiology, 73(10), 3196–3204. 10.1128/AEM.02610-06

Callahan, B. J., McMurdie, P. J., Rosen, M. J., Han, A. W., Johnson, A. J. A., & Holmes, S. P. (2016). DADA2: High-resolution sample inference from Illumina amplicon data. Nature Methods, 13(7), 581–583. 10.1038/nmeth.3869

Caporaso, J. G., Lauber, C. L., Walters, W. A., Berg-Lyons, D., Huntley, J., Fierer, N., Owens, S. M., Betley, J., Fraser, L., Bauer, M., Gormley, N., Gilbert, J. A., Smith, G., & Knight, R. (2012). Ultra-high-throughput microbial community analysis on the Illumina HiSeq and MiSeq platforms. The ISME Journal, 6(8), 1621–1624. 10.1038/ismej.2012.8

Crossette, E., Gumm, J., Langenfeld, K., Raskin, L., Duhaime, M., & Wigginton, K. (2021). Metagenomic Quantification of Genes with Internal Standards. mBio, 12(1), e03173–20. 10.1128/mBio.03173-20

Fierer, N., Jackson, J. A., Vilgalys, R., & Jackson, R. B. (2005). Assessment of soil microbial community structure by use of taxon-specific quantitative PCR assays. Applied and Environmental Microbiology, 71(7), 4117–4120. 10.1128/AEM.71.7.4117-4120.2005

Hayer, M., Wymore, A. S., Hungate, B. A., Schwartz, E., Koch, B. J., & Marks, J. C. (2022). Microbes on decomposing litter in streams: Entering on the leaf or colonizing in the water? The ISME Journal, 16(3), 717–725. 10.1038/s41396-021-01114-6

Hungate, B. A., Marks, J. C., Power, M. E., & Schwartz, E. (2021). The Functional Significance of Bacterial Predators. 12(2).

Hungate, B. A., Mau, R. L., Schwartz, E., Gregory Caporaso, J., Dijkstra, P., van Gestel, N., Koch, B. J., Liu, C. M., McHugh, T. A., Marks, J. C., Morrissey, E. M., & Price, L. B. (2015). Quantitative microbial ecology through stable isotope probing. Applied and Environmental Microbiology, 81(21), 7570–7581. 10.1128/AEM.02280-15

Meselson, M., Stahl, F. W., & Vinograd, J. (1957). EQUILIBRIUM SEDIMENTATION OF MACROMOLECULES IN DENSITY GRADIENTS. Proceedings of the National Academy of Sciences, 43(7), 581–588. 10.1073/pnas.43.7.581

Neufeld, J. D., Dumont, M. G., Vohra, J., & Murrell, J. C. (2007). Methodological considerations for the use of stable isotope probing in microbial ecology. Microbial Ecology, 53(3), 435–442. 10.1007/s00248-006-9125-x

Neufeld, J. D., Vohra, J., Dumont, M. G., Lueders, T., Manefield, M., Friedrich, M. W., & Murrell, J. C. (2007). DNA stable-isotope probing. Nature Protocols, 2(4), 860–866. 10.1038/nprot.2007.109

Nuccio, E. E., Blazewicz, S. J., Lafler, M., Campbell, A. N., Kakouridis, A., Kimbrel, J. A., Wollard, J., Vyshenska, D., Riley, R., Tomatsu, A., Hestrin, R., Malmstrom, R. R., Firestone, M., & Pett-Ridge, J. (2022). HT-SIP: A semi-automated stable isotope probing pipeline identifies cross-kingdom interactions in the hyphosphere of arbuscular mycorrhizal fungi. Microbiome, 10(1), 199. 10.1186/s40168-022-01391-z

Panijpan, B. (1977). The buoyant density of DNA and the G + C content. Journal of Chemical Education, 54(3), 172. 10.1021/ed054p172

Parada, A. E., Needham, D. M., & Fuhrman, J. A. (2016). Every base matters: Assessing small subunit RRNA primers for marine microbiomes with mock communities, time series and global field samples. Environmental Microbiology, 18(5), 1403–1414. 10.1111/1462-2920.13023

Radajewski, S., Ineson, P., Parekh, N. R., & Murrell, J. C. (2000). Stable-isotope probing as a tool in microbial ecology. Nature, 403(6770), 646–649. 10.1038/35001054

Satinsky, B. M., Gifford, S. M., Crump, B. C., & Moran, M. A. (2013). Use of Internal Standards for Quantitative Metatranscriptome and Metagenome Analysis. In Methods in Enzymology (Vol. 531, pp. 237–250). Elsevier. 10.1016/B978-0-12-407863-5.00012-5

Schwartz, E. (2007). Characterization of growing microorganisms in soil by stable isotope probing with H218O. Applied and Environmental Microbiology, 73(8), 2541–2546. 10.1128/AEM.02021-06

Simpson, A., Wood Charlson, E. M., Smith, M., Beilsmith, K., Koch, B., Walls, R. L., & Wilhelm, R. C. (2023). *A data standard for the reuse and reproducibility of any stable isotope probing-derived nucleic acid sequence (MISIP)* [Preprint]. Bioinformatics. 10.1101/2023.07.13.548835

Smets, W., Leff, J. W., Bradford, M. A., McCulley, R. L., Lebeer, S., & Fierer, N. (2016). A method for simultaneous measurement of soil bacterial abundances and community composition via 16S rRNA gene sequencing. Soil Biology and Biochemistry, 96, 145–151. 10.1016/j.soilbio.2016.02.003

Vyshenska, D., Sampara, P., Singh, K., Tomatsu, A., Kauffman, W. B., Nuccio, E. E., Blazewicz, S. J., Pett-Ridge, J., Louie, K. B., Varghese, N., Kellom, M., Clum, A., Riley, R., Roux, S., Eloe-Fadrosh, E. A., Ziels, R. M., & Malmstrom, R. R. (2023). A standardized quantitative analysis strategy for stable isotope probing metagenomics. *mSystems*, e01280–22. 10.1128/msystems.01280-22

